# ADAM-tRNA-seq: An Optimized Approach for Demultiplexing and Enhanced Hierarchal Mapping in Direct tRNA Sequencing

**DOI:** 10.1101/2025.05.19.654926

**Authors:** Rodrigo Alarcon, Daniel Köster, Stine Behrmann, Zoya Ignatova

## Abstract

Transfer RNAs (tRNAs) play an essential role in protein synthesis and cellular homeostasis, with their dysregulation associated with various human pathologies. Recent advances in direct RNA sequencing by the Nanopore platform have enabled simultaneous profiling of tRNA abundance, modifications, and aminoacylation status. However, the high sequence similarity among tRNAs and the lack of robust demultiplexing strategies reduce the accuracy and limit the scalability of current approaches. Here, we developed ADAM-tRNA-seq, a framework that addresses two key limitations of the Nanopore-based direct tRNA sequencing. First, we develop an RNA-based barcode demultiplexing method, employing a barcode embedded within the sequencing adapter that is recognized by the Dorado basecaller. Second, we designed a hierarchy-based mapping strategy that mitigates read loss due to multimapping by classifying reads at the isodecoder, isoacceptor, or isotype levels, thereby enhancing quantification accuracy. We validated ADAM-tRNA-seq using both synthetic tRNAs and a complex human tRNA pool, and systematically optimized it to achieve up to 99% classification precision. Together, these developments enable more accurate, scalable, and comprehensive characterization of tRNA pools in diverse sample types.

## INTRODUCTION

Transfer RNAs (tRNAs) are central components of the translation machinery with a canonical function in decoding sense codons of mRNA. The concentration and composition of cellular tRNA pools (or tRNAomes) differ between cell types, states, and tissues, and thus, play a crucial role in maintaining cell- and tissue-specific homeostasis and environmental stress response (1–10). Furthermore, several diseases are causally linked to dysfunctional tRNAs, resulting from genetic mutations in tRNA-encoding genes or auxiliary proteins involved in tRNA biogenesis, modifications, and aminoacylation (reviewed in (11–20)). Recently, inspired by developments in RNA therapeutics, innovative applications of tRNAs have emerged as potential therapies for a wide range of previously untreatable human conditions (20– 22).

The fundamental role of tRNAs in tissue and cell homeostasis has driven the development of various quantitative approaches to determine composition, expression levels, and modification patterns of tRNAs. tRNA-tailored microarrays were fundamental in the early stage of tRNAomics (5, 10). They can discriminate between tRNA species that differ by more than 8nt, and thus, well capture differences between most isoacceptors, but are limited in resolution to differentiate single isodecoders (5, 10, 23– 25). On the other hand, next-generation sequencing (NGS)-based approaches have achieved higher throughput and higher capacity and can reach isodecoder resolution (4, 26–32). These methods rely on reverse transcription and PCR amplification, whereby many tRNA modifications interfere with cDNA synthesis and may introduce bias in the libraries. Improved reverse transcriptases have enhanced the end-to-end processivity and efficiency of NGS tRNA-seq approaches by bypassing modifications (1, 9, 32, 33).

Nanopore technology supports direct RNA sequencing (d-RNA-seq) without reverse transcription. Recent developments in library preparation and data analysis have enabled direct tRNA sequencing (d-tRNA-seq) and broadened the scope towards quantifying tRNAomes, identifying modified nucleotides, and measuring tRNA aminoacylation levels (2, 34–38). However, some limitations still need to be addressed to fully unleash the potential of d-tRNA-seq as a benchmark technology to quantify the dynamics of cellular tRNA pools. The high sequence similarity between tRNA isodecoders, particularly in tRNAomes enriched with isodecoders (e.g., of higher eukaryotes), results in multimapping of sequencing reads and creates tRNA-specific biases in abundance estimation, which depends on the similarity and number of isodecoders per tRNA family.

Despite the demand, robust demultiplexing solutions for parallel sequencing of multiple samples are currently lacking. Previously developed strategies for demultiplexing, such as the community-driven DeePlexiCon (39), are now obsolete due to the discontinuation of the SQK-RNA002 chemistry, which has been replaced by the SQK-RNA004 kit. The implementation of this SQK-RNA004 chemistry has greatly improved the sequencing depth (i.e., gigabases generated per sequencing run), opening the door for upscaling and developing effective demultiplexing solutions. Recently developed methods, SeqTagger (40) and WarpDemuX (41), use barcode sequences in DNA adapters to classify reads by directly processing the raw signals from the Nanopore device. These strategies are able to demultiplex sequencing of both long and short RNAs. However, these methods require specifically trained models for the different sample types, sequencing chemistry, and barcode sequences, limiting their applicability and adaptability. As a result, d-tRNA-seq remains a complex, costly, and time-intensive method, limiting its scalability and broader applications.

Here we address these limitations by establishing ADAM-tRNA-seq (<u>A</u>dapter-based <u>D</u>emultiplexing <u>a</u>nd hierarchical <u>M</u>apping for <u>tRNA seq</u>uencing), an approach that uses a new hierarchy-based mapping strategy combined with barcodes integrated in the RNA adapters for efficient demultiplexing and read classification. All available isodecoder sequences of each tRNA family are used as reference. The mapping classifies reads to tRNA isotype, isoacceptor, or isodecoder, irrespective of the multimapping. This mapping approach reduced tRNA-specific biases and enhanced overall mapping depth. We also incorporated 16-nucleotide (nt)-long barcodes in the RNA adapters used in the preparation of tRNA libraries. This approach is readily compatible with existing tRNA library preparation protocols and data processing pipelines, and enables downstream analysis with minimal computational demand. We used *in vitro*-transcribed tRNAs (IVT-tRNAs) to optimize the demultiplexing parameters and validated the stringency of demultiplexing and mapping by profiling the tRNAome of human HEK293 cells.

## MATERIAL AND METHODS

### *In vitro* tRNA production

tRNAs were transcribed *in vitro* using the T7 transcription system as described previously (42), using two partially overlapping DNA oligonucleotides carrying the tRNA sequence and an upstream T7 promoter (Supplementary Table 1). Briefly, 25 μM of both oligonucleotides dissolved in 20 mM Tris-HCl (pH 7.5) were denatured for 2 min at 95 °C, annealed for 3 min at room temperature, followed by extension using 4 U/μL RevertAid Reverse Transcriptase (Thermo Fisher Scientific) for 40 min at 42 °C with 0.4 mM dNTPs. The purified dsDNA template was *in vitro* transcribed using 0.12 µg/µl T7 RNA polymerase overnight at 37 °C in the presence of 2 mM NTPs, 5 mM GMP, in 40 mM Tris-HCl pH 7.9, containing 0.01 mM Triton X-100, 3 mM spermidine, 10 mM MgCl_2_, 10 mM DTT. tRNAs were gel-purified and eluted in 50 mM KOAc (pH 7.0), containing 200 mM KCl, further purified by ethanol precipitation, and resuspended in DEPC-treated water. tRNA integrity was estimated by denaturing polyacrylamide gel electrophoresis.

### Cell culture and tRNA purification

HEK293 cells were grown in Dulbecco’s Modified Essential Medium (DMEM, Pan Biotech) supplemented with 10% fetal bovine serum (FBS, Pan Biotech), 1% penicillin/streptomycin (Gibco), and 1x GlutaMAX (Gibco) at 37 °C in 5% CO_2_. At 80% confluency, total RNA was isolated using TRIzol (Invitrogen) as described by the manufacturer. Total RNA was deaminoacylated by incubating in 100 mM Tris-HCl (pH 9.0) at 30 °C for 30 min. tRNAs were enriched using the RNA Clean & Concentrator (Zymo) following the manufacturer’s instructions to separate the RNAs with a size between 17 and 200 nt. RNA quality was determined using the Bioanalyzer RNA 6000 Nano (Agilent).

### tRNA library preparation

The library preparation was performed by adapting the Nano-tRNAseq protocol (2) using the SQK-RNA004 kit (ONT) as described below. 5’ and 3’ barcoded RNA adapters (Supplementary Table 2) were mixed to a concentration of 2.7 µM in annealing buffer (10 mM Tris-HCl, pH 7.5, containing 50 mM NaCl), and annealed RNA adapters were ligated to the *in vitro-*transcribed or cellular tRNA samples at molar ratio of 1.2:1 at 21 °C for 2 h with 1 µL of T4 RNA Ligase 2 (10 U/µL) (NEB), and 0.5 µL of RiboLock RNase Inhibitor (40 U/μL) (Thermo Scientific) in T4 RNA Ligase 2 Buffer and 15% PEG 8000 (NEB) (43). tRNA was purified using AMPure RNAClean XP beads (Beckman Coulter) and eluted in nuclease-free water.

DNA oligos matching the RTA adapters (ONT) were annealed as described above for the RNA adapters, and ligated to 200 ng of barcoded tRNA at a molar ratio of 2:1 for 30 minutes at 21 °C in NEBNext Quick Ligation Reaction Buffer (NEB), 1.5 µL of T4 DNA ligase (2000 U/µL) (NEB), and 0.5 µL of RiboLock RNase Inhibitor (40 U/μL) (Thermo Scientific) in a final volume of 15 µL. Reverse transcription of the adapted tRNAs was performed by adding 13 µl of nuclease-free water to the ligation reaction, 2 µl of 10 mM dNTPs, 8 µl of 5× Maxima H Minus Reverse Transcriptase Buffer, and 2 µl of Maxima H Minus Reverse Transcriptase (Thermo Scientific). The mixture was incubated at 60 °C for 1 h, followed by 85 °C for 5 min. The samples were then purified with 50 µL AMPure RNAClean XP beads (Beckman Coulter) and eluted in nuclease-free water.

The final adapter ligation was carried out by adding 5 µL of RLA (RNA Ligation Adapter), 8 µL of 5x NEBNext Quick Ligation Reaction Buffer (NEB), and 5 µL of T4 DNA ligase (2000 U/µL) (NEB, M0202M). The mixture was incubated for 20 min at 21 °C, purified with AMPure RNAClean XP (Beckman Coulter) beads, and eluted with REB (Elution Buffer).

### Nanopore sequencing and basecalling

Multiplexed libraries were pooled to an equimolar concentration in a volume of 12 µL, mixed with 25.5 µL of LIS (Library Solution) and 37.5 µL of SB (Sequencing Buffer). The samples were then loaded into a MinION RNA flow cell (FLO-MIN004RA) following the ONT SQK-RNA004 protocol. Libraries were run in a MinION instrument using the MinKNOW software version 24.11.10 for 40 to 72 hours, selecting POD5 files as outputs. The data was processed using the Dorado basecaller version 0.9.1 in a super accurate mode version 5.1.0 (rna004_130bps_sup@v5.1.0).

### Demultiplexing

Basecalled reads were first filtered to retain those with a length between 50 and 200 nt. Barcode classification was performed using the Dorado software (v0.9.1), employing a custom barcode classification module. Due to the incomplete sequencing of the 5’ barcode, only the 3’ barcode was considered for classification using the “rear_only_barcodes = true” parameter. Multiple sets of scoring parameter sets were evaluated using *in vitro*-transcribed tRNAs to assess the classification accuracy. The parameter set providing the highest accuracy was then selected and applied to the cellular tRNA dataset for the final barcode classification.

### Benchmarking parameters

To evaluate the performance of the demultiplexing strategy, we used IVT-tRNAs and considered two primary benchmarking metrics: precision and recovery. These metrics were calculated for each barcode and subsequently averaged to yield an overall performance estimate for each set of demultiplexing parameters. Precision was defined as the proportion of correctly classified tRNA reads for a specific barcode, relative to the total number of reads for that tRNA across all barcodes, calculated as:

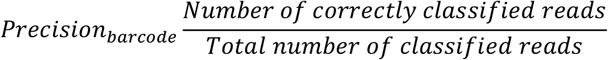

Equation 1: Calculation of the precision per barcode to evaluate the proportion of correctly classified reads of one tRNA.

Recovery was defined as the proportion of tRNA reads for a given barcode that were correctly classified, relative to the total number of reads derived from that tRNA as determined independently of demultiplexing. To estimate the total number of reads for each tRNA, sequencing reads were first mapped directly to the reference sequences without applying any demultiplexing. Recovery per barcode was calculated as:

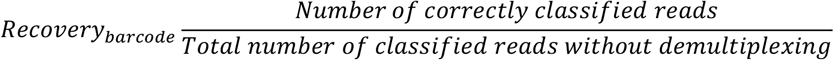

Equation 2: Calculation of the recovery per barcode to estimate the proportion of reads that are correctly classified relative to the total number of reads for that tRNA without demultiplexing.

### Mapping of human cellular tRNAome

The reference dataset for human tRNAs was generated from the GtRNAdb (GRCh38) (44), selecting entries from the “High Confidence tRNA Gene Set” with an overall tRNA model score above 60. Separate reference files were created for each isoacceptor family to enable hierarchical mapping. Demultiplexed reads were aligned independently to each reference file using the Burrows-Wheeler Aligner (BWA-MEM) with parameters optimized for tRNA reads (-W 13 -k 6 -x ont2d -T 10) (2). For each read, the resulting alignments were compared across all isoacceptors. Based on the alignment scores, reads were hierarchically classified as uniquely mapped to a specific isodecoder, isoacceptor, or isotype. This classification was used to retain reads at the highest possible level while minimizing multimapping ambiguity.

## RESULTS

### RNA-based demultiplexing workflow

Previous approaches developed for tRNA demultiplexing use barcodes incorporated within the DNA adapters that cannot be basecalled by the standard Nanopore software (39–41). We reasoned that embedding barcodes within the RNA adaptors would be a more practical solution for tRNA libraries, allowing for barcode basecalling using the established RNA basecallers. For this, we modified a 16-nt long stretch of the RNA adapter sequences originally designed for Nano-tRNAseq (2), while retaining the first two and last six nt of the 5’ adapter to preserve ligation efficiency and avoid interference with adapter recognition (Figure 1A). Additionally, we kept the GC content consistent, avoided homopolymeric stretches of three or more identical nucleotides, and adjusted the complementary 3’ adapter sequences accordingly. To ensure robust demultiplexing, we designed four unique barcodes with a minimum pairwise edit distance of 11 nucleotides (Supplementary Table 2).

**Figure 1.**
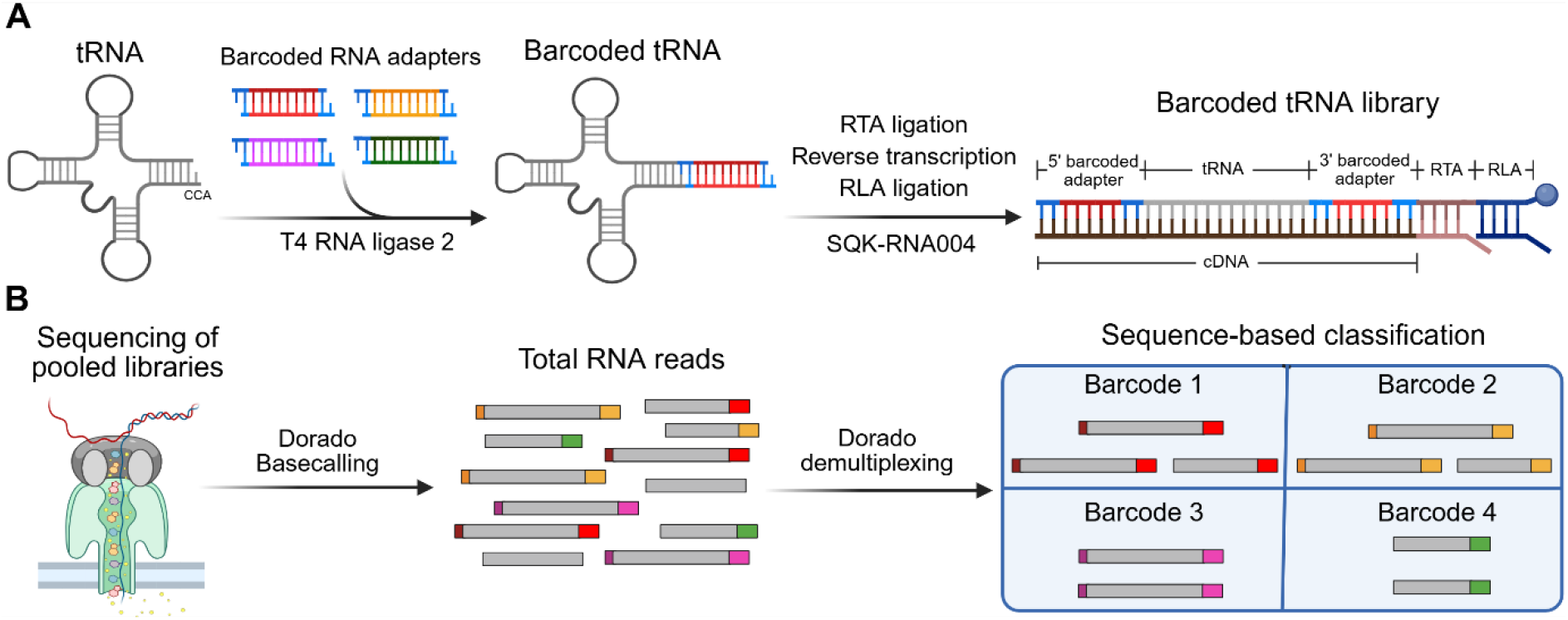
Scheme of the RNA-based barcoding and demultiplexing used in the ADAM-tRNA-seq. (**A**) Library preparation using four RNA barcodes (color-coded). (**B**) Sequencing and demultiplexing of tRNAs. Total basecalled reads are represented with color-coded adapters at left (5’ adapter) and right (3’ adapter) ends of the reads. Detected reads without 5’ or any adapter are also depicted.

To test our demultiplexing strategy, we used the barcoded RNA adapters to generate libraries with four IVT-tRNAs originating from different isotypes (Figure 1A, Supplementary Table 1). The libraries were pooled, sequenced on a MinION device, and basecalled using the Dorado RNA model “<u>sup@v5.1.0</u>”. We then performed barcode-based classification using the Dorado Demux model (ONT), a tool originally developed for demultiplexing of Nanopore DNA-seq (Figure 1B). During analysis, we consistently observed incomplete sequencing of the 5’ RNA adapters, likely due to technical limitations inherent to the Nanopore platform. Thus, we only considered the 3’ adapter barcode in the demultiplexing procedure by setting the built-in option rear_only_barcodes to “true”. Finally, the demultiplexed reads were mapped using BWA-MEM.

### Demultiplexing parameter optimization

A notable feature of the Dorado Demux tool is its ability to demultiplex reads using custom barcodes and custom scoring parameters. To achieve the best performance when demultiplexing tRNA reads, we systematically tested various scoring parameters individually to evaluate their impact on classification performance by calculating demultiplexing precision and recovery (Supplementary Figure 1). We observed that “midstrand_flank_score” and “min_flank_score” only have a minor influence on precision. Hence, we did not consider them in the further optimization. We then proceeded to iterate combinations of `max_barcode_penalty`, `min_barcode_penalty_dist`, `min_separation_only_dist`, `flank_left_pad`, and `flank_right_pad`. The parameter intervals were chosen based on the individual parameter tests’ results, considering the length and the edit distance of our customized barcodes. Our raw data indicated that certain reads did not fully capture the entire poly-A tail at the end of the 3’ barcode. Thus, we decided against incorporating the complete tail in our right flank parameter. We calculated demultiplexing precision and read recovery rate across all sequencing data sets, leading to a total evaluation of 648 parameter combinations (Supplementary Table 3). We reached a demultiplexing precision of up to 99% (Figure 2A).

**Figure 2.**
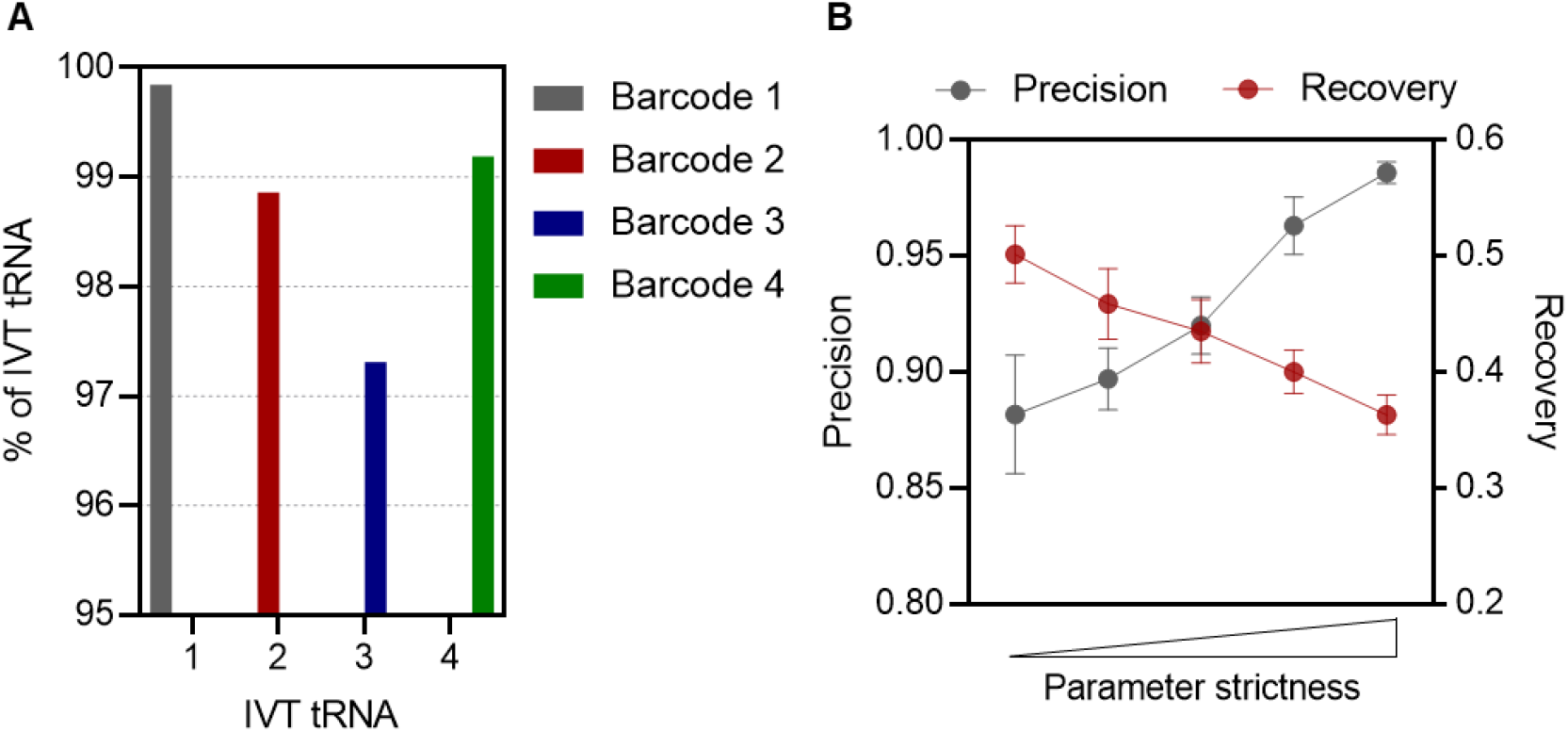
IVT-tRNA demultiplexing and optimization. (**A**) Percentage of each IVT-tRNA classified per barcode. The four distinct IVT-tRNAs were labelled with barcodes 1, 2, 3, and 4. (**B**) Demultiplexing performance with five parameter sets with different strictnesses (Supplementary Table 4). Values are means ± SD (n = 3 independent experiments).

We observed that the demultiplexing performance was highly dependent on the demultiplexing parameters. With increased strictness of the demultiplexing, the precision increased, while the recovery decreased. From this analysis, we chose five sets of parameters that achieved precision levels ranging from 90% to 99% while exhibiting the highest recovery rates (Figure 2B and Supplementary Table 4). This systematic optimization enabled us to determine conditions that maximized read recovery while maintaining a defined level of demultiplexing precision. As the recovery is inversely proportional to the precision, the user should define the demultiplexing parameters based on the desired precision to retain the maximum number of reads.

### Hierarchy-based mapping

Cellular tRNAs exhibit high similarity between isodecoders, representing a unique mapping challenge for d-tRNA-seq analysis. In addition, the average sequencing accuracy for Nanopore d-tRNA-seq is still below 99%. Moreover, tRNAs are pervasively modified, which creates basecalling errors (2, 34). This makes it challenging to accurately distinguish between reads originating from different isodecoders or isoacceptors with high similarity, resulting in some reads mapping to multiple reference sequences. Previous approaches have addressed this issue by selecting a single isodecoder sequence for each isoacceptor (2) or including all isodecoder sequences in a single reference file (34). However, these approaches can lead to an underestimation of uniquely mapped reads or to extensive multimapping, which results in a loss of a significant proportion of reads. Multimapped reads, namely those that map with high quality to multiple reference sequences, are discarded by default by the mapping tools. Identifying how to treat multimapping reads is crucial in d-tRNA-seq and depends on the downstream analysis. When comparing the abundance between different isodecoders, only reads that can unambiguously be identified as a specific isodecoder should be kept, while multimapped reads aligning to multiple isodecoders with the same mapping quality should be discarded. In contrast, if a downstream analysis is focused on the distribution of isoacceptors, reads that multimap to different isodecoders of the same isoacceptor family can be kept and pooled. Similarly, when analysing isotype distributions, all isoacceptors for a given isotype can be grouped, thus allowing for retention of reads that multimap to distinct isoacceptors within the same isotype family, while discarding those that map to different tRNA isotypes with equal mapping quality (Figure 3A).

**Figure 3:**
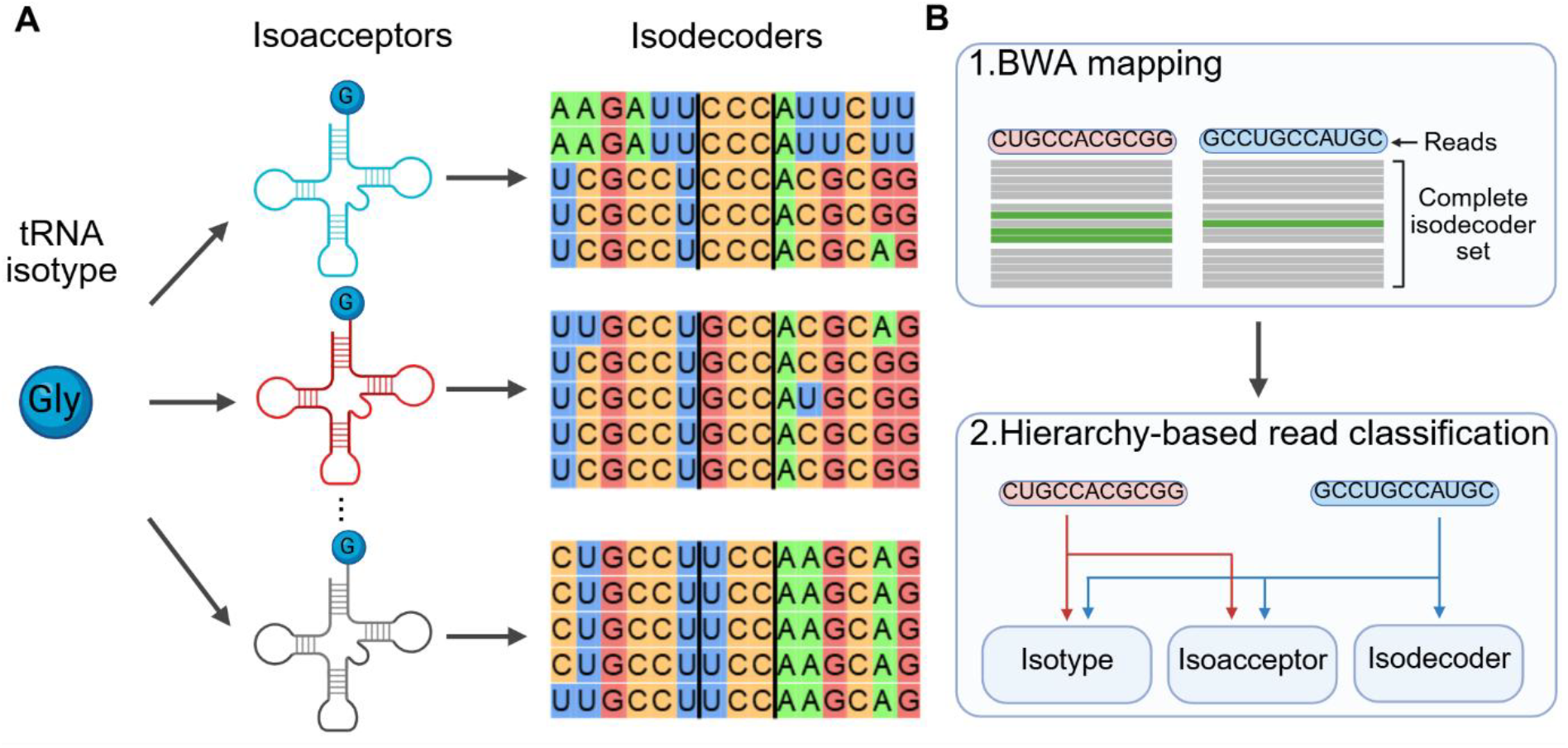
Hierarchy-based read classification for mapping reads used in ADAM-tRNA-seq. (**A**) Hierarchical separation of tRNA types used to classify mapped reads. Each nucleotide is color-coded, and the tRNA anticodons are displayed between two black lines. (**B**) Mapping strategy. The pipeline consists of two steps: 1. BWA mapping and 2. hierarchy-based read classification.

Considering all this, we developed a workflow – named “hierarchy-based read classification” – that minimizes read losses at different levels of hierarchy, i.e., isodecoder, isoacceptor, and isotype. Instead of using a single reference file, our mapping pipeline groups all the isodecoder sequences by their isoacceptor and aligns each read to each group independently. Following mapping, multimapping reads are not discarded. Instead, we select the highest alignment quality per read and assign it to a specific isodecoder if possible. If the alignment score is equal for more than one isodecoder reference within the same group, the read is assigned to the corresponding isoacceptor. Consecutively, if the alignment score is equal across different isoacceptor groups within the same tRNA isotype, the read is assigned to that isotype (Figure 3B). Thus, the individual mapping strategy can operate in three modes based on the user’s needs, providing isodecoder, isoacceptor, or isotype resolution.

To assess the mapping performance with the hierarchy-based read classification, we performed d-tRNA-seq of total tRNA isolated from HEK293 cells. We mapped the basecalled reads using BWA-MEM, analyzed the mapping quality score for each isodecoder, and classified the reads at the three different levels (i.e., tRNA isotype, isoacceptor, and isodecoder). We observed a wide distribution of the mapping quality scores between tRNA isoacceptors (Supplementary Figure 2). To capture highly modified tRNAs that would result in low mapping quality, we set a minimum mapping quality threshold of 10. Then, we compared the hierarchy-based mapping with other mapping algorithms (2, 34). Our mapping strategy uses comparable computational demand relative to simpler alignment schemes (Supplementary Table 5). Using our hierarchy-based classification approach, we observed a notable increase in the percentage of mapped reads identified at all levels of resolution compared to the other two mapping strategies (Figure 4A). The percentage of classified mapped reads increased significantly for all isotypes (Figure 4B) and most isoacceptors (Supplementary Figure 3).

**Figure 4:**
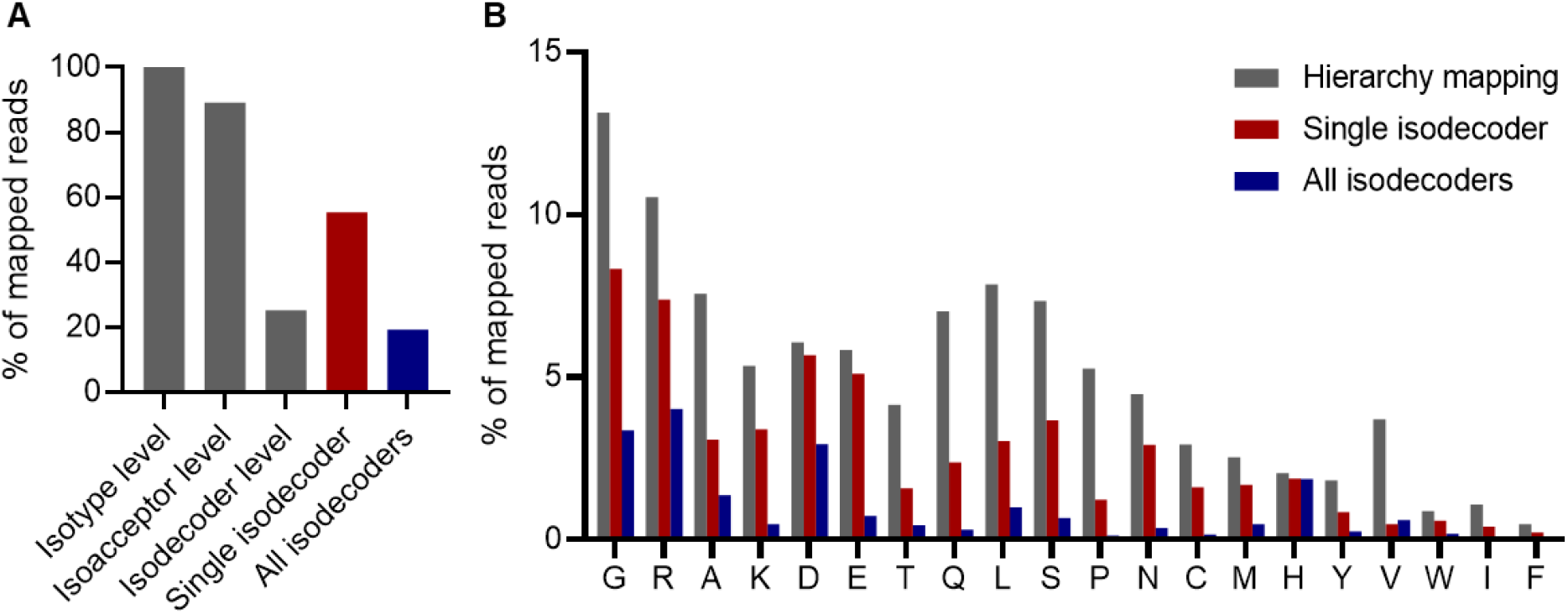
Hierarchy-based classification improves sequencing depth and reduces multimapping bias. **(A)** Percentage of total mapped reads that are classified using hierarchy-based mapping (grey), or mapped to single isodecoders (2) (red), or all isodecoder sequences in a single reference file (34) (blue). **(B)** Percentage of mapped reads grouped by isotype using the three mapping strategies as in panel A. tRNA isotypes are designated with the single-letter code of the cognate amino acid.

At the isodecoder level, our mapping strategy performs comparably to the strategy that uses all isodecoder sequences in a single reference (Figure 4A), as both methods consider all isodecoders without allowing multimapping. However, we observed an increase in uniquely mapped reads using the hierarchy-based method. This is likely due to its ability to avoid sequence similarities across isoacceptor groups that otherwise would elevate the likelihood of multimapping. At the isoacceptor and isotype levels, the hierarchy-based mapping recovers substantially more reads than mapping with all isodecoders as reference (Figure 4B and Supplementary Figure 3). Furthermore, our strategy outperforms the approach using a single representative isodecoder per isoacceptor, which inherently limits its resolution to the isoacceptor and isotype levels (Figure 4A). Notably, mapping at the isoacceptor level, our approach successfully recovered reads that were previously discarded due to poor mapping quality against the representative isodecoder reference (Supplementary Figure 3). This improvement in read recovery was not uniform across all isotypes and isoacceptors. These differences likely stem from variations in isodecoder copy number and reflect differences in isodecoder sequence similarity within each tRNA family.

### Evaluation of cellular tRNAome

To assess the performance of ADAM-tRNA-seq in a complex biological tRNA data set, we isolated the total tRNA from HEK293 cells. We split the sample into four aliquots. Three of them were spiked with different synthetic IVT-tRNAs. Each sample was uniquely barcoded using one of the four RNA adapter barcodes during library preparation. For barcode 1, no spike-in was added, and this sequencing was used as a baseline mapping control to evaluate potential misassignment from the barcodes. Equal molar amounts of all four tRNA libraries were pooled and sequenced. Reads were basecalled and demultiplexed as described above using the highest accuracy parameters set and mapped using hierarchy-based read classification at all three levels (Supplementary Tables 6, 7, and 8).

We first analyzed the distribution of the spike-in tRNA reads across barcodes to assess the demultiplexing precision. Each spike-in tRNA was detected in its corresponding library, with a precision higher than 99% (Figure 5A). Comparison between the four samples reveals that the abundance of mapped reads to tRNA isotypes remains consistent across the four barcodes, with a standard deviation below 1% for all isotypes. Together, these data indicate a uniform read classification among barcodes without bias (Figure 5B) and show that ADAM-tRNA-seq can be used to sequence multiple cellular tRNA samples with high reproducibility and precision.

**Figure 5:**
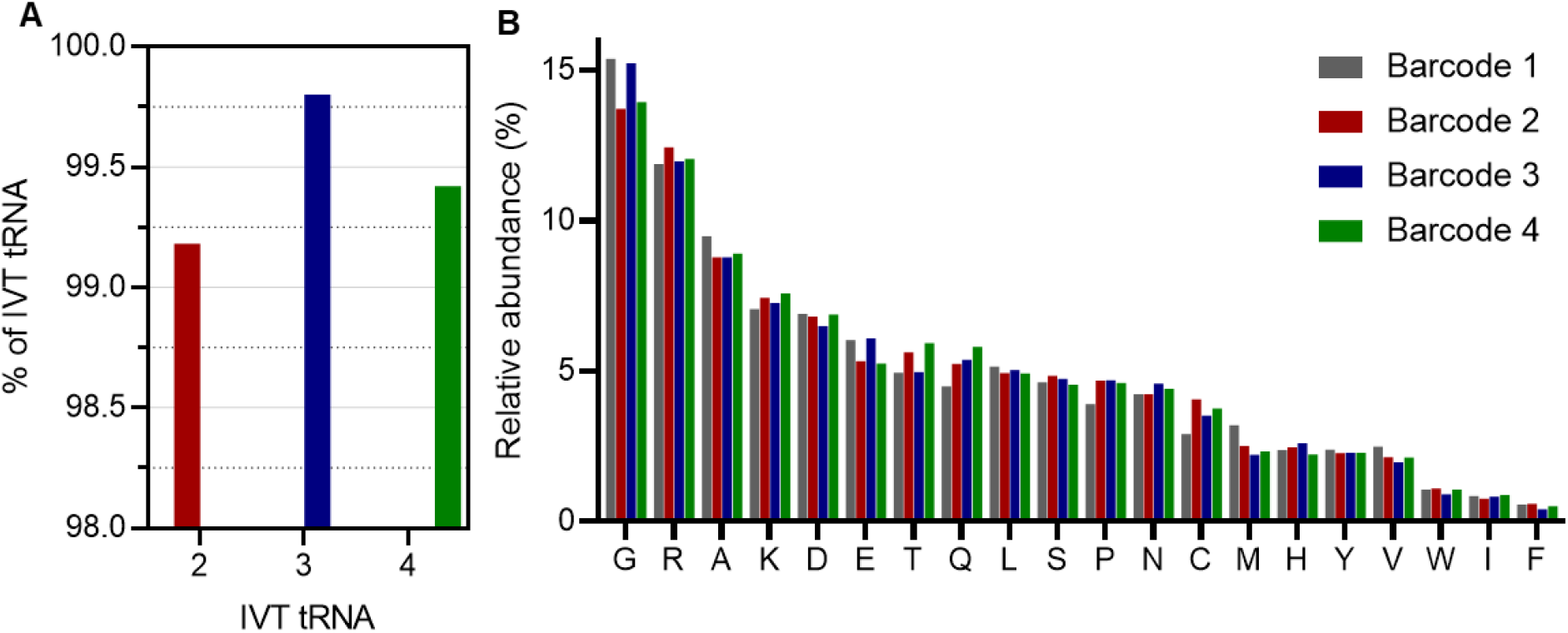
ADAM-RNA-seq of the tRNA pool of HEK293 cells. (**A**) Percentage of spike-in IVT-tRNA per barcode. Library with barcode 1 was not spiked. IVT-tRNAs were used as spike-ins for the libraries with barcodes 2, 3, and 4 (the same as in Figure 2). (**B**) Relative abundance of each tRNA isotype. tRNA isotypes are designated with the single-letter code of the cognate amino acid.

## DISCUSSION

In this study, we present ADAM-tRNA-seq, including a new RNA-based demultiplexing and mapping strategy optimized for d-tRNA-seq using the Nanopore platform. Our approach enhances classification precision and read assignment by integrating the barcode detection with a hierarchical mapping framework. Relative to the currently simpler alignment schemes, the hierarchical mapping significantly improves read retention and mapping accuracy. The current implementation uses the BWA-MEM algorithm, although the mapping strategy is aligner-independent. Alternative alignment tools, e.g., minimap2 (45) or Parasail (46), could offer improvements in speed or even accuracy for specific read types.

A key advantage of ADAM-tRNA-seq is the use of barcodes embedded in the RNA adapters, which can be recognized by RNA basecallers, thus enabling native demultiplexing of tRNA reads without relying on DNA-derived signals or basecalling models trained on DNA data (40, 41). This makes the method inherently compatible with changes in Nanopore sequencing chemistry and basecalling models, which would otherwise require retraining of DNA demultiplexing tools. Moreover, while demonstrated here with tRNAs, the RNA barcode strategy can be expanded to other RNAs, provided the library preparation includes RNA adapters. This allows for potential integration into broader RNA sequencing workflows, including those targeting small RNAs, viral RNAs, or structured noncoding RNAs. We also develop a hierarchical mapping approach that classifies reads at the level of tRNA isodecoder, isoacceptor, or isotype based on mapping quality and sequence identity. This allows users to adjust the resolution of the analysis according to the experimental goals. Importantly, this mapping framework increases the proportion of retained reads compared to traditional strategies (2, 34) that either exclude multimapping reads or use minimal reference sets. Our approach resolves ambiguities through the hierarchical classification, enhancing mapping depth and reducing quantification bias resulting from tRNA isodecoders’ sequence similarity.

ADAM-tRNA-seq multiplexing capability can be expanded by incorporating more barcodes into the workflow. The performance of the new barcodes should be benchmarked to ensure that the demultiplexing accuracy is maintained. In the barcode design, the GC content among the used barcodes should be kept consistent, and the flanking adapter sequences, which determine ligation efficiency, should remain unchanged. The performance of the demultiplexing may be improved by increasing the barcode length. In addition, the RNA barcodes can be combined with signal-based demultiplexing approaches to further increase the number of samples measured in parallel. Lastly, while the method was evaluated on human tRNAs, it is designed to be broadly applicable to tRNAomes from other species.

In summary, ADAM-tRNA-seq provides a robust and flexible solution for accurate tRNA profiling. It enhances both precision and data retention, supports multiplexing, and can accommodate different experimental needs, thus facilitating the scalability and efficiency of d-tRNA-seq and d-RNA-seq in general.

