## Supplementary Figures for "ADAM-tRNA-seq: An Optimized Approach for Demultiplexing and Enhanced Hierarchal Mapping in Direct tRNA Sequencing"

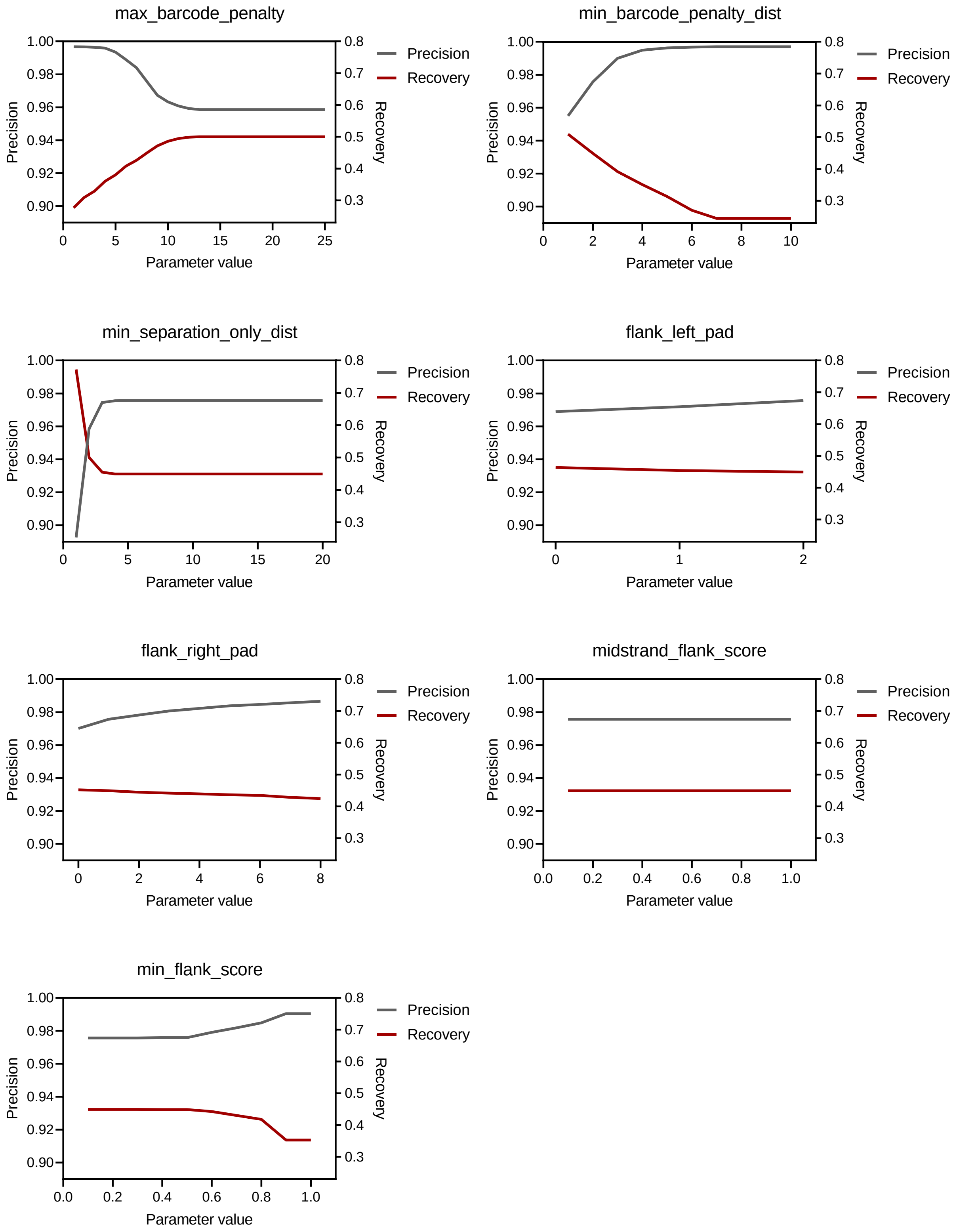


**Supplementary Figure 1:** Effect of the individual demultiplexing parameters on precision and recovery using Dorado Demux for sequencing of IVT tRNAs. The evaluated parameter was changed as indicated, while the other parameters were kept constant at the following default values: max_barcode_penalty = 8, barcode_end_proximity = 100, min_barcode_penalty_dist = 2, min_separation_only_dist = 6, flank_left_pad = 2, flank_right_pad = 1, front_barcode_window = 175, rear_barcode_window = 175, midstrand_flank_score = 0.1, min_flank_score = 0.1.

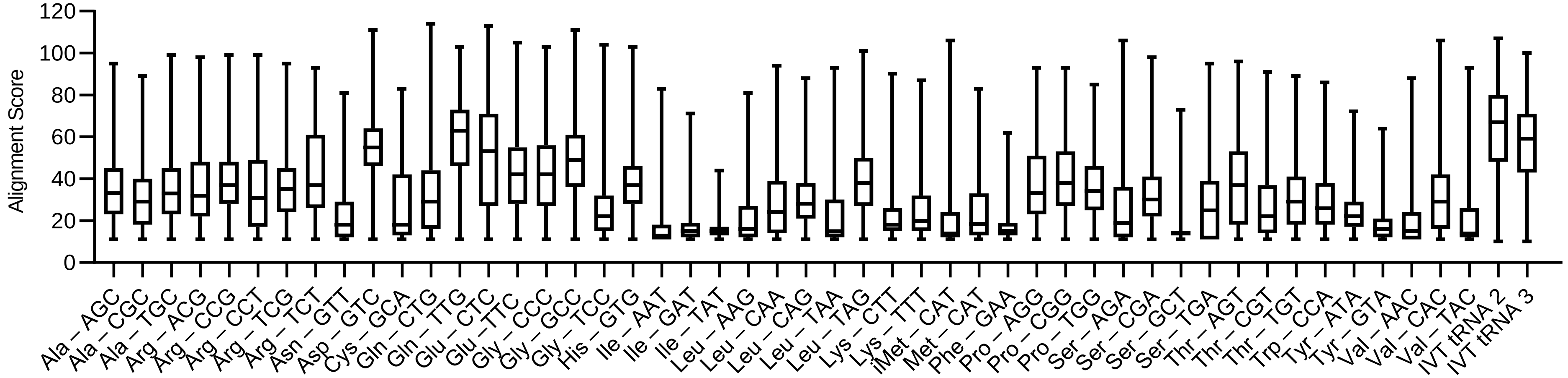


**Supplementary Figure 2:** Alignment score for each tRNA isotype. Total tRNA isolated from HEK293 cells were sequenced and processed using ADEM-tRNA. Reads with an alignment score below 10 were discarded. IVT tRNAs 2 and 3 are identical to those used in Fig. 2.

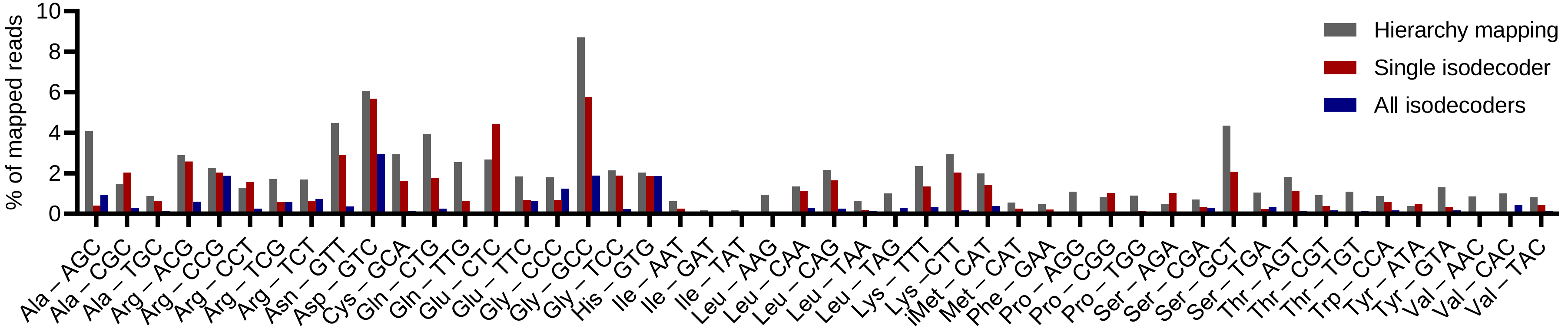


**Supplementary Figure 3:** Percentage of total mapped reads classified to a single isoacceptor using hierarchy-based mapping, mapping with single isodecoders, and mapping with all isodecoder references.
